# A Guide for Active Learning in Synergistic Drug Discovery

**DOI:** 10.1101/2024.09.13.612819

**Authors:** Shuhui Wang, Alexandre Allauzen, Philippe Nghe, Vaitea Opuu

**Affiliations:** Laboratoire de Biophysique et Evolution, UMR CNRS-ESPCI 8231 Chimie Biologie Innovation, PSL University, Paris, France; LAMSADE, Universite Paris-Dauphine, PSL University, Paris, France

## Abstract

Synergistic drug combination screening is a promising strategy in drug discovery, but it involves navigating a costly and complex search space. While AI, particularly deep learning, has advanced synergy predictions, its effectiveness is limited by the low occurrence of synergistic drug pairs. Active learning, which integrates experimental testing into the learning process, has been proposed to address this challenge. In this work, we explore the key components of active learning to provide recommendations for its implementation. We find that molecular encoding has a limited impact on performance, while the cellular environment features significantly enhance predictions. Additionally, active learning can discover 60% of synergistic drug pairs with only exploring 10% of combinatorial space. The synergy yield ratio is observed to be even higher with smaller batch sizes, where dynamic tuning of the exploration-exploitation strategy can further enhance performance.

## 1 Introduction

Single-drug treatment targeting cancer often leads to drug resistance, i.e. by upregulation of enzymes involved in alternative signaling pathways [1]. For example, Chemotherapy using Cisplatin, a widely used drug in various cancers by producing DNA damage, triggers the overexpression of GSTP1 and reduces drug efficacy [2]. For this reason, chemotherapy is often based on combinations of two already approved drugs. Combination treatments are based on synergistic drug pairs, where the combined effect of the two drugs exceeds the sum of their individual effects. For instance, synergy is expected to arise when perturbing metabolic or regulatory pathways that compensate for each other, so both need to be broken to yield a significant effect [3]. As a tendency, the proportion of single-drug therapy has decreased by about 40%, being substituted by drug combination therapies [4].

However, identifying synergistic drug combinations requires exploring a large space of conditions. Drugcomb [5] is a meta-database that aggregates 34 distinct campaigns, comprising 8397 drugs, 2320 cell lines, and 739964 drug combinations. Except for the large combinatorial space, synergy is a rare phenomenon. Oneil [6] and Almanac [7] are the most studied datasets, displaying 3.55% and 1.47% of synergistic drug pairs, respectively. The rise of high-throughput drug combination screening enables the generation of large datasets. However, conducting an exhaustive search for identifying highly synergistic pairs is time-consuming and expensive [8]. The Oneil data set was produced using experimental automated platforms, performing 25 rounds of 896 measurements, leading to 22,737 conditions in total. The ALMANAC campaign conducted 304,549 experiments with a platform over 340 rounds. Due to the high costs of these extensive studies, it is generally infeasible for a typical biological laboratory to conduct such large-scale campaigns. Therefore, it is crucial to rapidly identify effective drug combinations despite the large combinatorial space and low discovery rate.

To increase the likelihood of discovering targets given limited resources, computational algorithms were developed to predict synergy based on publicly available data. For instance, the DREAM dataset, composed of 118 drugs and 85 cell lines, has been provided as a learning set to develop artificial intelligence approaches to synergy discovery [9]. These methods offer biologists a preliminary evaluation tool to assess the potential value of conducting experimental studies. Although traditional machine learning (ML) algorithms such as regression and decision trees were largely used to predict the effect of drug combinations, Baptista et al. showed that deep learning (DL) algorithms now outperform them [10]. For example, DeepSynergy [11] is a DL algorithm composed of a single multi-layer perceptron (MLP) that predicts synergy using chemical and genomic descriptors as inputs. Larger neural networks are now leveraging the wealth of chemical databases (i.e., Chembl) to improve the prediction performances, which is directly inspired by the leaps observed in natural language translation and text generation [12]. As the representation used as input of the algorithm seems crucial to reaching high performances, DeepDDS [13] exploits the topology of molecules by representing them as graphs whereas GAECDS [14] uses graphs to represent drug-drug relationships. Torkamannia et al. [15] showed that incorporating protein-protein interactions (PPI) improves by 2% the prediction accuracy over algorithms without PPI, showing the importance of considering the cellular context. To exploit even more massive DL architectures, [16, 17] exploits more complicated architectures by including transformers.

However, these approaches focus on improving the prediction of synergy by using different drug and cell features. Berti et al. [18] are the first to combine a computational method with an active learning framework, to guide wet-lab experiments for the screening of synergy pairs. In the framework of active learning, instead of predicting which M measurements to perform in vitro using an AI algorithm, it divides the measurements by *k* batches to perform sequentially using the data obtained all along dynamically. Active learning was shown to improve by ∼ 5-10 times the detection of highly synergistic drug combinations compared with randomly choosing the combinations. RECOVER is composed of two layers: the 1) is an MLP that creates a numerical representation of molecules from the Morgan fingerprint [19], and 2) is a second MLP taking two molecules representations to predict the Bliss synergy score [20]. RECOVER is first pre-trained with the Oneil dataset [6], then it is used to select and measure batches of drug combinations iteratively. Along the procedure, the batches produced up to that point are used to refine sequentially RECOVER parameters. Similar strategies are being used in drug discovery [21, 22], biological network optimization [23] and bioengineering [24, 25].

Despite the successful application of active learning, several questions remain unclear. How to compose an active learning framework for the detection of synergistic drug combinations? As shown in Fig. 1, active learning comprises available data, an AI algorithm to evaluate new samples, and selection criteria for prioritizing the evaluation of drug pairs. We investigated here the elements composing an active learning framework, which we finally benchmarked against simulated scenarios of drug synergy campaigns. The first component is the AI algorithm used to select drug combinations for experimental measurement. To identify the most suitable algorithm, we conducted a benchmarking process focused on data efficiency, as these algorithms often function in low-data environments. Furthermore, we evaluated their generalization capability across new cell types and drugs, which confirmed the inherent difficulty of these tasks [18]. The selection process driven by the AI algorithm relies on the extrapolation-exploitation trade-off. We investigated several simulated experimental campaigns, showing the striking importance of the batch size. We show that 1488 measurements scheduled with active learning allowed us to recover 60% (300 out of 500) synergistic combinations, saving 82% experimental time and materials (without using any strategy, it requires 8253 measurements to obtain 300 synergistic combinations).

**Figure 1.**
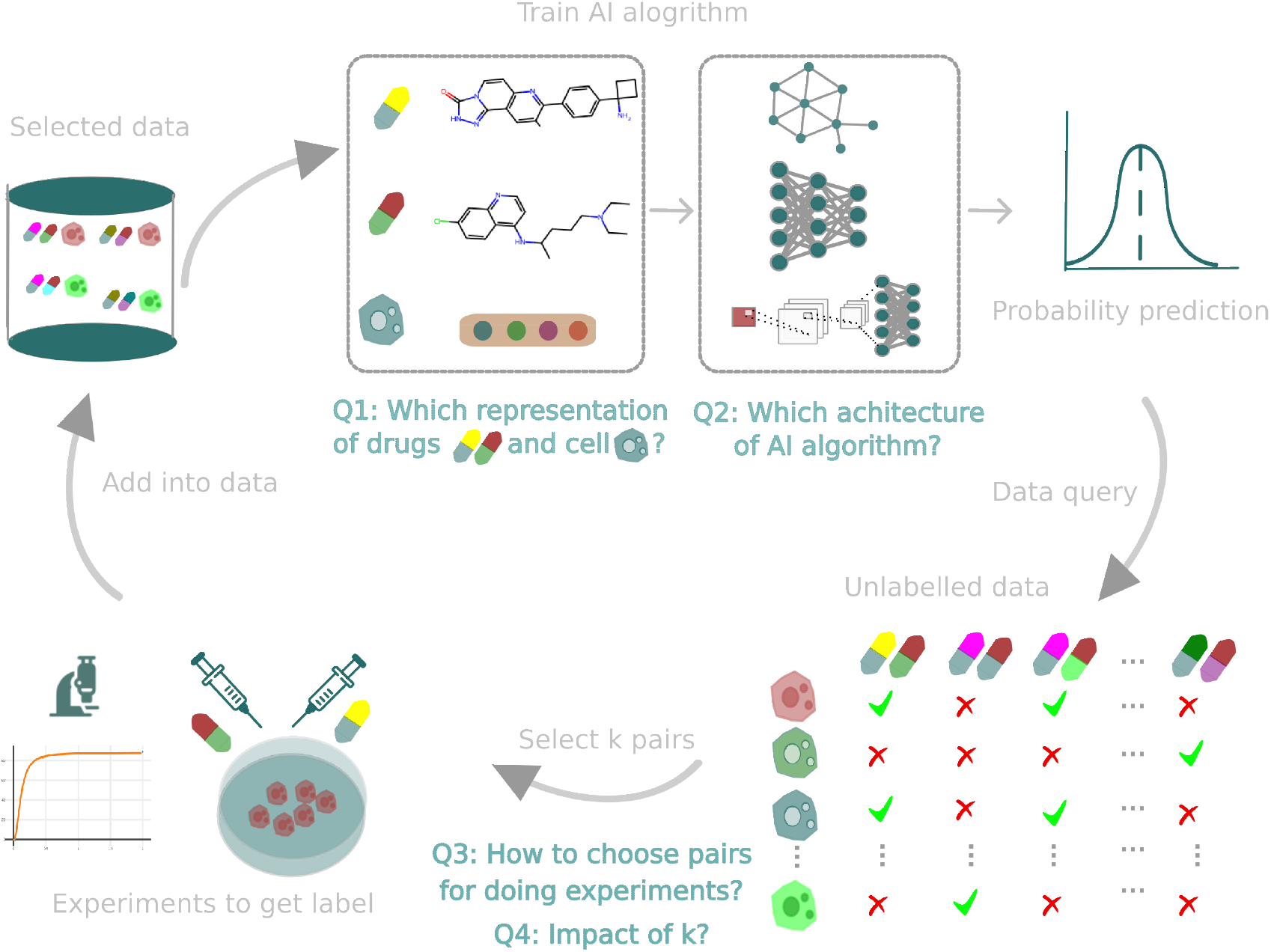
Active Learning cycle in synergistic drug discovery. The diagram illustrates the key components of the active learning workflow. Starting with an initial training dataset of drug combinations, the AI model learns from the data and suggests new candidates (hits) for experimental evaluation. These new results are then incorporated into the training set. The selection strategy involves choosing kk samples from a pool of drug combinations with unknown synergy. The strategy can be either explorative—aiming to improve model accuracy by selecting diverse samples—or exploitive, targeting combinations with a high likelihood of synergy. Our focus is on determining the optimal AI algorithm by addressing (Q1) feature selection and (Q2) algorithm architecture. Additionally, we explore (Q3) the best selection strategy and (Q4) the appropriate batch size for experimental validation.

## 2 Results

### 2.1 Data-efficient algorithms

Active learning requires an AI algorithm capable of learning starting from a small amount of training data. Thus, we benchmarked the three components for AI-based synergy prediction in data efficiency: i) the AI algorithms, ii) the molecular features describing molecules, and iii) the features describing the targeted cell.

For this benchmark, we used the Oneil [6] datasets, where we defined the drug pairs that yielded an experimental LOEWE synergy score greater than 10 as synergistic [26]. The Oneil dataset is composed of 15,117 measurements, compromising 38 drugs and 29 cell lines (corresponding to 74.15% of all combinations) including 3.55% synergistic drug pairs. Then, we used the precision-recall area under curve (PR-AUC) score to quantify the detection of synergistic drug pairs. In each testing case, the model was trained five times.

To make a prediction, AI algorithms take numerical representations of cells and drugs as input referred to here as features. To evaluate the approach in a low data regime, we used 10% of the data as validation and randomly selected 10% data from the remaining data as a training dataset. First, we benchmarked 5 distinct molecular features: OneHot encoding representation, Morgan fingerprints [19], MinHashed atompair fingerprint up to a diameter of four bonds (MAP4) [27], Molecular access system (MACCS) [28], and the representation pre-trained from ChemBerta2 [12]. To test these, we used the MLP AI algorithm along with the gene expression as cellular features. Using the MLP allows us to test molecular feature combinations used in synergy predictions by NN. Besides, we compared three permutation invariant combination operations: Sum, Max, and Bilinear operations from RECOVER (see details in methods). However, neither the molecular representation nor the combination operation displayed any striking gain in prediction quality, as shown in Fig. 2a. However, Morgan fingerprint with the addition operation displayed the highest prediction performance, significantly better than the OneHot (with two-sampled t-test p-value=0.04).

**Figure 2.**
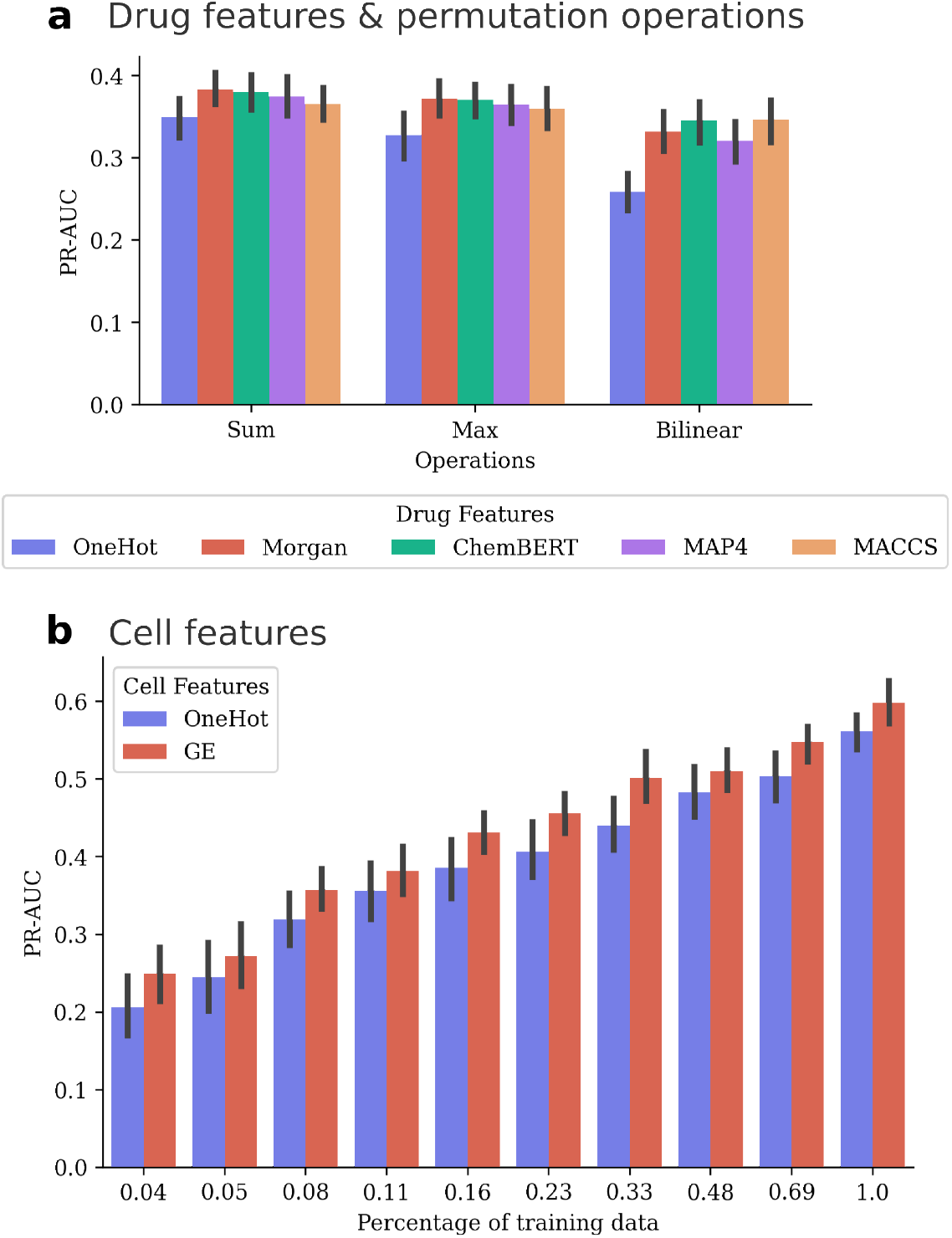
Retrospcetive testing shows the effect of drug features, invariant permutation operations and cell features. (a) Comparison of performance using different drug representation features—ChemBERT, Morgan fingerprints, and OneHot encoding—in combination with various permutation operations including Sum, Max, and Bilinear. (b) Comparison of performance using different cell representations, specifically OneHot encoding and gene expression data from the GDSC (Genomics of Drug Sensitivity in Cancer) database. The MLP model uses Morgan fingerprints as drug features and additive operation.

The cellular features allow the algorithm to account for the cellular environment of the targeted cell for making predictions. For cellular features, we compared a trained representation and the genetic single-cell expression profiles, obtained from the Genomics of Drug Sensitivity in Cancer (GDSC) database [29]. To test them, we used the MLP, the Morgan fingerprint molecular feature, and the addition combination. In this scenario, we used 10% of the data as validation and various percentages of data as a training dataset. Our results showed that using the single-cell expression profiles significantly improved the prediction quality, it achieved 0.02-0.06 gain in PR-AUC (with p-value=0.05) (Fig. 2b). Moreover, this result remained consistent as we varied the training set size.

Next, we compared AI algorithms ranging from parameter-light to -heavy. For the parameter-light algorithm, we tested the logistic regression (LR) and XGBoost gradient boosting (XGBoost) algorithms, which are typical ML algorithms used in drug property prediction [30]. In parameter-medium, we tested a neural network (NN) algorithm composed of 3 layers of 64 hidden neurons trained using the backpropagation algorithm. In parameter-heavy, we tested three algorithms based on deep learning, where DeepDDS GCN and DeepDDS GAT use the molecular topology to encode the drug representation, whereas DTSyn uses a transformer architecture [16] to extract molecular information. The number of parameters for these algorithms ranges from 700k (NN) to 81M (DTSyn).

We then tested the algorithms for data efficiency by evaluating their performances across increasing training sizes, where we used the Morgan fingerprint and gene expression profile. We observed two learning phases according to the performance trend shown in Fig. 3a. In the first phase (*<*48% of the training set), algorithms displayed substantial gains in learning performances as the training size increased. In the second phase (*>* 48% of the training set), the performance of algorithms displayed lower performance gains, where the MLP gain in performance is on average ∼ 0.07 between 6000 and 12000 data points. The latter means that all algorithms essentially reached the maximum potential from the 6000 data; therefore, adding measurements beyond this limit is not strongly beneficial for the training.

**Figure 3.**
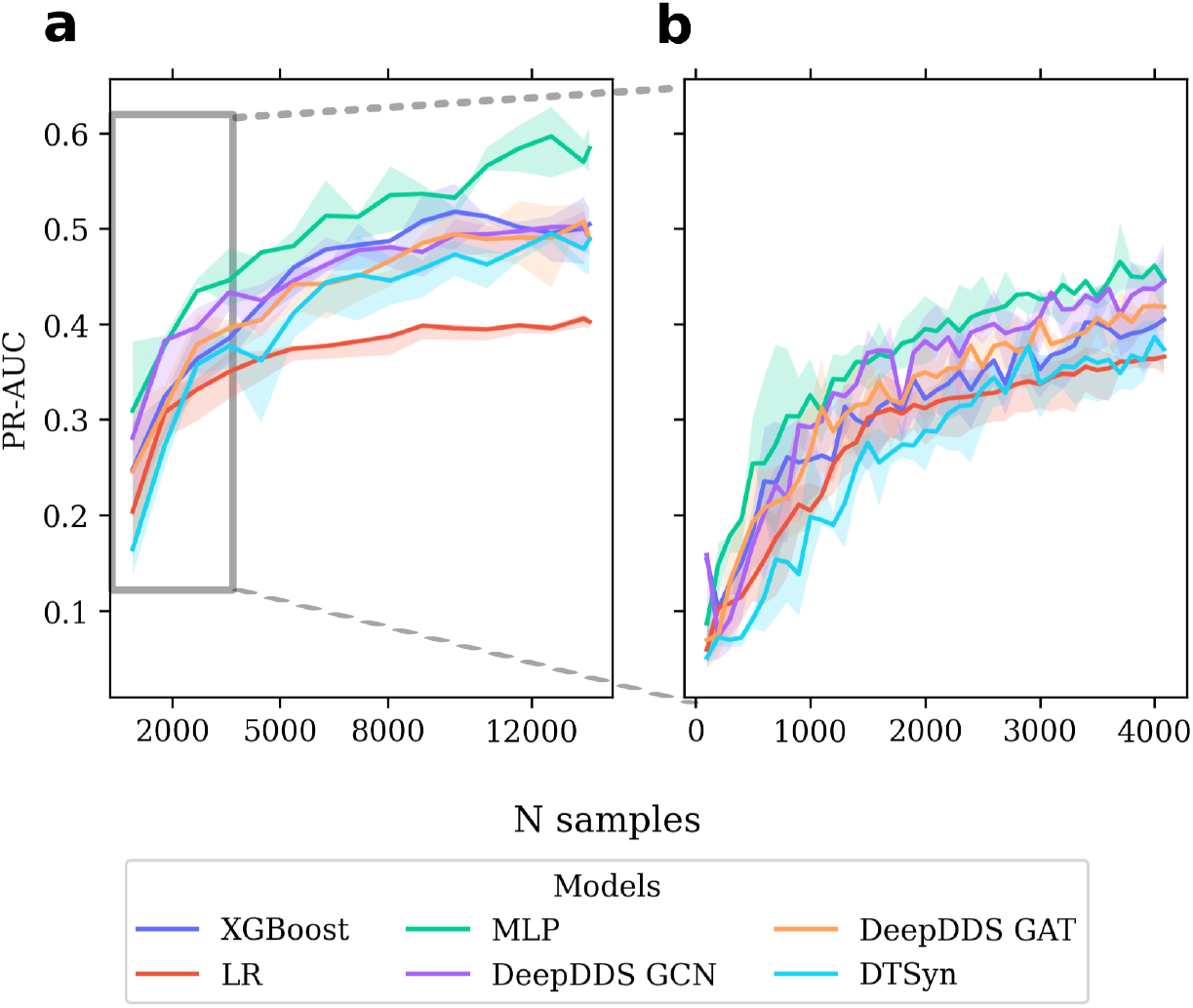
Performance of algorithms with varying proportions of training data. The X-axis shows the number of training data points, and the Y-axis displays the PR-AUC on the validation dataset. Solid lines represent average performance over 10 random cross-validations, with shaded areas indicating the 0.9 quantile range. (a) Performance of MLP and other algorithms across different training dataset sizes. (b) Zoomed view of algorithm performance in the low-data regime.

The highest performance (averaged over 10 replica PR-AUC=0.60) is achieved by the MLP algorithm at 88% (∼12,000 data points) in the plateau phase. All the others performed similarly (PR-AUC=0.49 on average at 88%) except for the LR which performed poorly at PR-AUC=0.40. In Almanac, we observed similar performance across models; however, except LR, these performances did not reach the plateau phase (see Fig. SI1).

Next, we examined the performances with a smaller increment (*k* = 100 data points) in the first phase, see Fig. 3b. These results showed that the highest performance gains happened with as few as 2000 data points (corresponding to 15% of the training set), where the average gain is 0.38. During this first phase, all methods, even LR, displayed similar performance gains; however, we observed that DTSyn, which is parameter-heavy, displayed the slowest performance curve. At least *>*15% of the training set was required to train DTSyn to reach the accuracy observed with the LR. In contrast, the average performance of the MLP is slightly higher than all other algorithms across the first phase.

Unexpectedly, our results mitigated the importance of molecular features as all of them performed similarly. Adding information about the topology of molecules or using pre-trained features learned on large molecular datasets did not provide any noticeable benefit. In contrast, the gene expression incorporating the cellular information significantly improved performances consistently across multiple training data sizes. This result emphasizes the unique effect of drugs in specific cell types. In general, our results showed that there are typically two learning phases: the first phase where the learning performance increases quickly, and the plateau phase where increasing the training size does not drastically improve the performance. Therefore, only measuring as few as 48% of all combinations is typically sufficient to obtain a predictive power of PR-AUC=0.51 with the MLP, which displayed the best performance overall. In the lowest data regime (*<*1000 training data points), our results suggest that even significantly large NN (2M parameters) can still be as predictive as the MLP or the LR. However, the most complex algorithm, DT-Syn (with 81 million parameters), demonstrated only marginal improvements (0.15), indicating that there is still potential to scale up the size of the MLP to further enhance its predictive performance if needed. The heavier algorithms are difficult to tune for specific applications, while parameter-light algorithms do not offer flexibility in usage. In contrast, the MLP does allow flexibility in the hyper-parametrization, which partially explains the performance difference observed here.

### 2.2 Testing generalization for novel drug and cell types

The greatest challenge in drug property prediction is ensuring that the algorithm generalizes beyond the drugs, cell lines, and tissues in the training set. However, it is not clear which type of AI algorithm can effectively extract general patterns across drugs or cell types. Consequently, we benchmarked seven AI algorithms for their generalization capabilities across drugs, cell types, and tissues (details in Materials and methods 6.5).

To perform the benchmark, we used the Oneil dataset that we partitioned as follows:

- Leave-combination-out: Data points were grouped by drug pairs across cell types, which were then partitioned into 80% training set and 20% test set, ensuring that drug combinations were not seen with any cell type during the training. We trained and then evaluated the average error on 50 random replicas of the partitioning.
- Leave-drug-out: Data points were grouped by drug, we then performed 20-fold cross-validation, where the data points corresponding to 1 or 2 drugs were kept in the test set of each fold whereas the others were used for training. On average, the test set is composed of 750 data points.
- Leave-cell-out: Data points were grouped by cell lines and partitioned such that the test set consisted of the data points corresponding to 1 cell line whereas the rest 28 cell lines were used for training. To evaluate models, we performed 29 cross-validation.
- Leave-tissue-out: Data points were grouped by tissue and performed a leave-one-out cross-validation where all the data points corresponding to one tissue were kept for validation whereas the other 5 were used for training.

The results are shown in Table 1 and Fig. 4a-d. In the leave-combination-out scenario, the average performance of the tested methods is PR-AUC = 0.52. As expected, LR performed poorly compared to more sophisticated approaches, which achieved an average PR-AUC of 0.36. XGBoost outperformed LR (p *<* 0.05), but was still significantly less accurate than neural network-based methods (p *<* 0.05). Among MLP, DeepDDS, and DTSyn, only marginal and non-significant differences were observed.

**Table 1.** Performance of each algorithm (PR-AUC value) on leave-combination-out, leave-drug-out, leave-cell-out and leave-tissue-out experiments. The results is *avg* ± *std*. The value in bold is to indicate the algorithm which achieved the highest mean PR-AUC.

| Algorithms | Combination | Drug | Cell | Tissue |
| --- | --- | --- | --- | --- |
| LR | 0.36±0.19 | 0.15±0.15 | 0.41±0.19 | 0.29±0.11 |
| XGBoost | 0.49±0.22 | 0.17±0.17 | 0.45±0.17 | 0.35±0.07 |
| MLP | 0.59±0.20 | **0.37±0.23** | 0.51±0.17 | 0.38±0.07 |
| DeepDDS GCN | 0.60±0.22 | 0.25±0.17 | 0.52±0.17 | 0.38±0.08 |
| DeepDDS GAT | **0.62±0.21** | 0.36±0.23 | **0.52±0.16** | **0.40±0.07** |
| DTSyn | 0.59±0.19 | 0.34±0.24 | 0.51±0.14 | 0.34±0.07 |

**Table 2.**
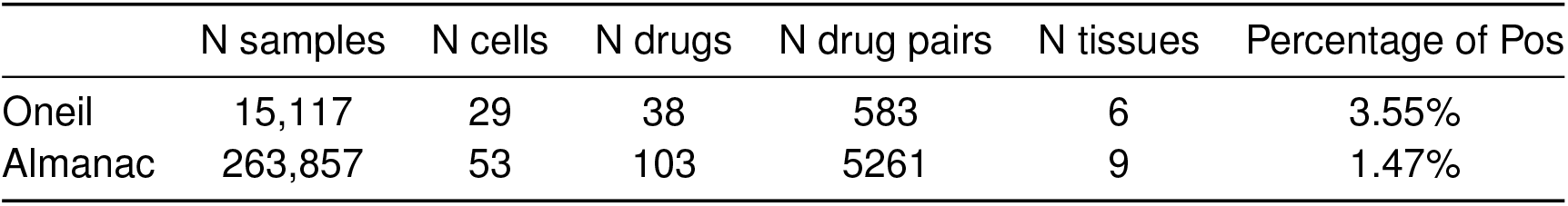
The summary of datasets, including the number of samples, the number of cells, the number of unique drug pairs, the number of tissues and percentage of synergies within each dataset.

**Table 3.** The hyperparameters setting for MLP model.

| Hyperparameters | Values |
| --- | --- |
| Hidden size for final synergy prediction | [ 128, 64] |
| Hidden size for dimension reduction for drugs | [256, 128] |
| Hidden size for dimension reduction for cell | [256, 128] |
| Dropout rate | 0.1 |
| Activation function | Relu |
| Learning rate | $10^{-4}$ |
| Maximum epoches | 100 |

**Table 4.** The summary of deep learning methods.

| models | N hyperparameters | Cell features (D) | Drug features (D) | Datasets |
| --- | --- | --- | --- | --- |
| MLP | 698, 113 | GDSC (908d) | ChemBert (1200d) | Oneil, Almanac, Dream |
| DeepDDS-GCN | 1,165,462 | CCLE(954d) | GCN (atoms 5d) | Oneil |
| DeepDDS-GAT | 2,817,538 | CCLE (954d) | GAT (atoms 5d) | Oneil |
| DTSyn | 80,920,706 | CCLE, LINCS (978d) | GCN ( atoms 5d) | Oneil, Almanac, Dream, FLOBAK, FORCINA and YOHE |

**Figure 4.**
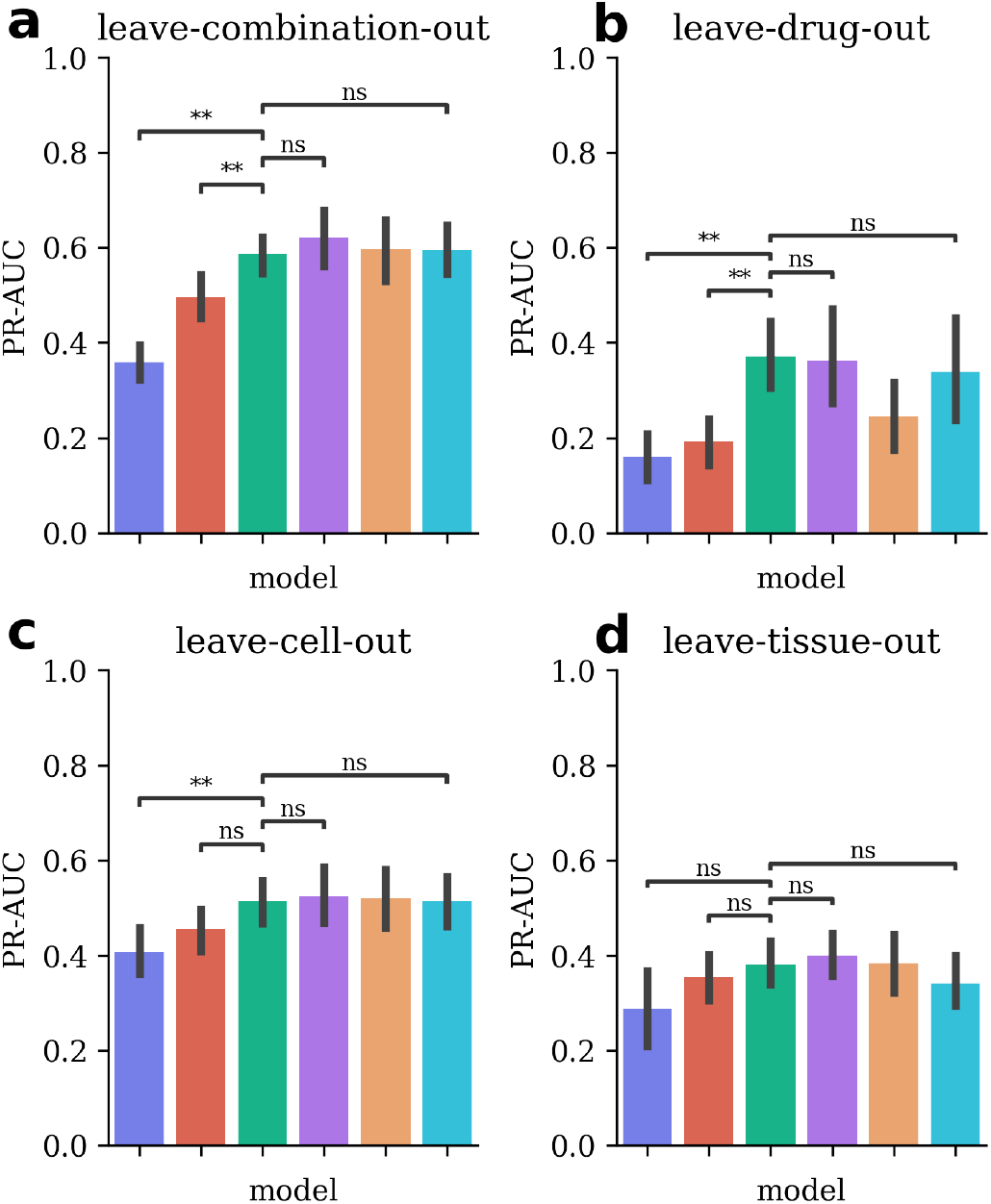
Performance of distinct models across different scenarios, illustrating their generalization abilities. Each scenario represents a different data-splitting approach, showcasing various aspects of algorithm performance. (a) Leave combination out: random split of data. (b) Leave drug out: 5% of drugs are excluded. (c) Leave cell out: All drug combinations for a specific cell are excluded as validation. (d) Leave tissue out: All cells from a specific tissue are excluded as validation. X-axis shows all models, then Y-axis is the PR-AUC value. ns: 0.05 *< p* ≤1, *: 0.01 *< p* ≤ 0.05, **: *p <*= 0.01. Performance is evaluated on each task by using cross validation.

For leave-drug-out, the average performance of the tested methods is PR-AUC = 0.26, significantly lower than predicting new drug combinations (p *<* 0.05). Similar to the earlier scenario, LR had the poorest performance with a PR-AUC of 0.15. However, in this partitioning, XGBoost performed poorly compared with the neural network-based algorithms, with an average PR-AUC of 0.19. MLP achieved the highest performance with an average PR-AUC of 0.37.

In the leave-cell-out scenario, the average performance across all methods is PR-AUC = 0.48, higher than predicting novel drugs (p *<* 0.05). Both LR and XGBoost had the worst performance with an average PR-AUC of 0.41 and 0.45, significantly lower than the neural network-based algorithms. MLP and other parameter-heavy algorithms performed comparably, with an average PR-AUC of 0.52.

In leave-tissue-out, the average performance across all methods is PR-AUC = 0.36, lower than in leave-cell-out (p *<* 0.05). Unlike earlier partitions, all methods performed similarly, except for LR, which had a PR-AUC of 0.29.

Our results indicate that when some data are already available for the drugs to be tested, NN algorithms can consistently detect new synergistic combinations. Besides, for the NN, on average, 43% of the true synergies in the test set were discovered. In contrast, when no information about a drug is present in the training set (leave-drug-out), the synergy prediction is poor, only 22% of true synergies can be discovered, see Fig. SI2b. This poses a significant challenge for synergy prediction—and molecular property prediction in general—because it suggests that information does not easily generalize to new drugs. Incorporating new cell types is more favorable for prediction, provided that similar cell types exist in the training set. Indeed, by removing all cell types from one tissue, we observed a drastic drop in performance. These results suggest that no AI algorithm can truly generalize beyond the training set on its own, highlighting the need for integrated approaches such as active learning.

### 2.3 Active learning to accelerate drug synergy screening

In the previous sections, we benchmarked input features and algorithms. In this section, we detailed the construction of an active learning workflow. Subsequently, we examined the effects of varying batch size and different sample selection strategies.

Active learning approach can dynamically select which drug combination to measure in each consecutive batch, and prioritize drug combinations with a high likelihood of synergy. The typical active learning workflow has two components: (a) the predictor AI algorithm that is learned from the available data to assign predicted synergy scores to drug pairs not yet tested, and (b) the selection strategy, based on the predicted scores, identifies *k* (batch size) drug combinations to be tested experimentally by a platform, which is then incorporated into the training data of the AI algorithm.

Here, estimating the uncertainty of predictions from AI algorithms is a key step for guiding the selection process. Typically, an ensemble approach is employed to determine the prediction confidence for each unmeasured pair [31]. In this approach, *M* instances of an AI algorithm are trained simultaneously. The average prediction is used as a synergy score *S* while the standard deviation of the predictions is used to quantify the uncertainty *σ*(*S*). Both quantities are then used to select the combination to measure next.

For the predictor, we chose the MLP because it performed the best in data efficiency and extrapolation across drugs, cell types, and tissues (see Sections 2.1-2.2). Therefore, we started our investigation with the MLP as a predictor, in combination with Morgan fingerprint molecular features and gene expression for cellular features.

For the selection strategy, there are three approaches. The first approach uses the *S* score to select potentially the most synergistic drug combinations. We refer to this strategy as the exploitation strategy. The second approach uses *σ*(*S*) to select the drug combinations that the algorithm struggles to predict, which corresponds to the most informative measurements to perform. We refer to this strategy as an exploration strategy. The third approach is a mixture of the two scores (*S, σ*(*S*)), which we refer to as a hybrid strategy.

For the experimental tests, the critical parameter is the batch size (the number of experimental tests per cycle), which is limited by the laboratory equipment capacity. For example, a semi-automatic platform runs with a batch size of *k* = 168 whereas a fully automatic platform reaches *k* = 896. Depending on this, the optimal active learning framework may also vary.

First, we evaluated the selection strategies by simulating two scenarios, as realistically as possible, using the Oneil dataset. In the first scenario, we simulated a manual platform, yielding *k* = 168 measurements per batch; in the second scenario, we simulated an automatic platform yielding *k* = 896 measurements (details are in Materials and methods 6.6). In our simulation, we screened *C* = 5 cell types as follows:

- Initialization: *k* random combinations are extracted from the Oneil dataset and used as the initial training dataset.
- Training: *k* instances of the AI algorithm are trained using the current training set.
- Criterion evaluation: We evaluate both criteria on all drug-drug-cell triplets which are not yet measured.
- Selection: We select the five cell types that yield the highest criterion, depending on the strategy. Then, for each cell, we select an array of *k/*5 drug combinations for each cell type which yields the highest criterion too.
- Experimental measurements: To simulate the experimental screening of a batch, we reveal the measurements corresponding to the five cell types and the selected drugs, which we add to the training data for the next iteration.

Then, we repeated all the steps from the training until all entries from the Oneil dataset were revealed. Our selection process mimics experimental constraints by manipulating only five cell types simultaneously [6]. To measure the quality of the model with the current training set, we extracted 10% of the remaining data as validation to compute the PR-AUC. We compared these active learning strategies to our null selection strategy (baseline), where the selection of training data points to improve the model is made randomly—not informed by the learning algorithm.

The results of our simulations showed that active learning significantly increased the synergistic pairs detection; for example, as few as 4 batches (4476 measurements) of exploitation strategy with the automatic platform allowed us to detect 50% of all synergistic pairs (500 pairs in total). When active learning is not used (baseline), at least 9 batches of measurements are required, corresponding to approximately 4 weeks of experimental labor. Consistently, active learning strategies significantly improved the performance of the learning algorithm, meaning that the training sample constructed are indeed more informative (Fig. 5a). Our simulations however did not yield significant differences between exploration strategy and exploitation strategy. We posit that because the number of samples that are indeed synergistic is so few, the highly synergistic ones also exhibit a high standard deviation for the predictions; indeed, the synergy is inversely proportional to standard deviation (Pearsonman *ρ* = 0.90) (Fig. SI3). Therefore, both strategies tend to select the same type of pairs. We validated further our results using the Almanac dataset (Fig. SI4), showing that the size of the combinatorial space did not change the trend of synergy detection.

**Figure 5.**
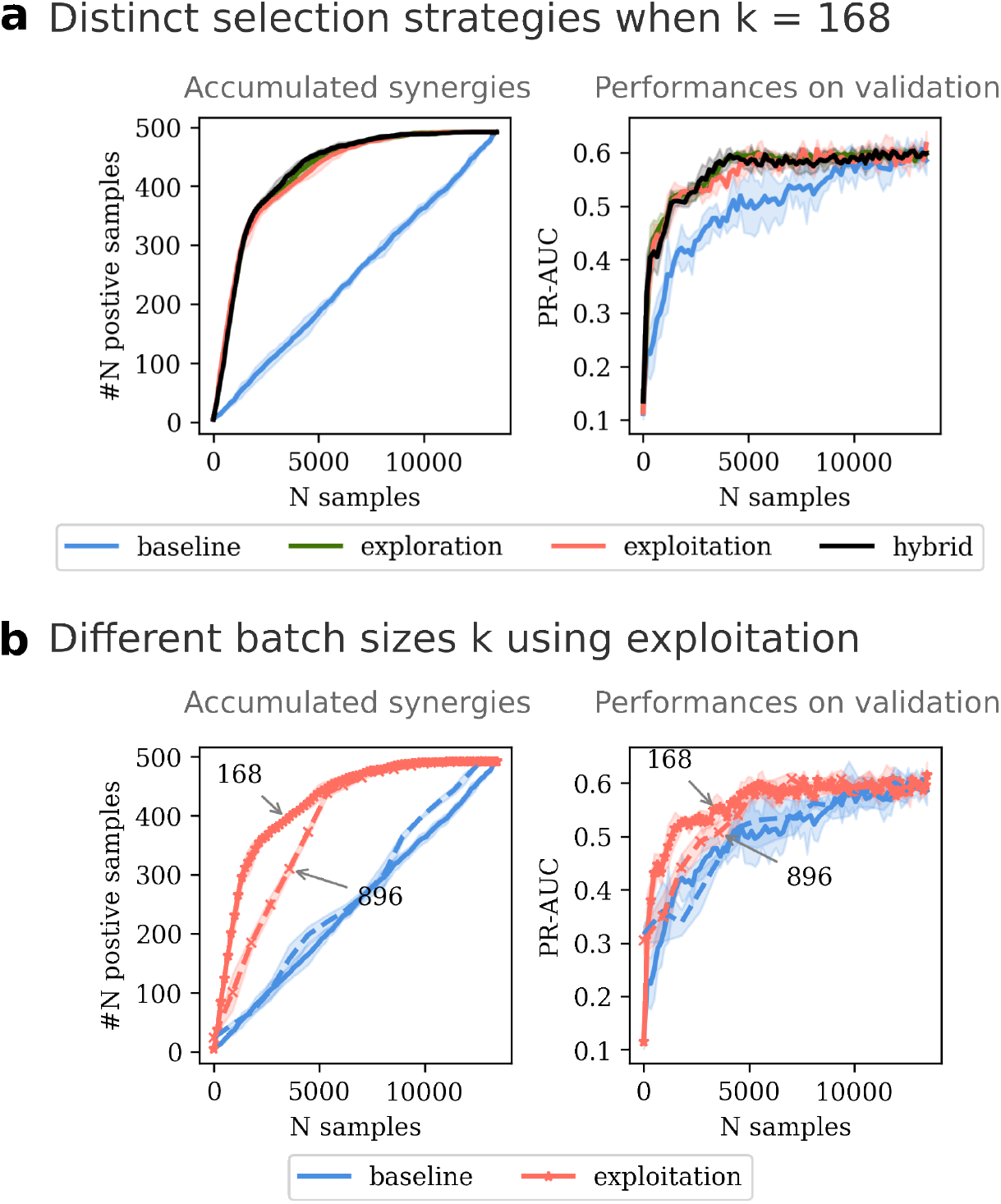
The impact of query selection strategy and batch size on synergy discovery and algorithm performance. Left Panels: Accumulated number of synergistic samples as experiments progress across different rounds. Right Panels: Performance of the NN algorithm as additional data is sequentially added. (a) Comparison of selection strategies—exploration (green), exploitation (pink), and hybrid (black)—with a batch size *k* = 168. (b) Comparison of outcomes with batch sizes *k* = 168 (solid lines) versus *k* = 896 (dashed lines).

**Figure 6.**
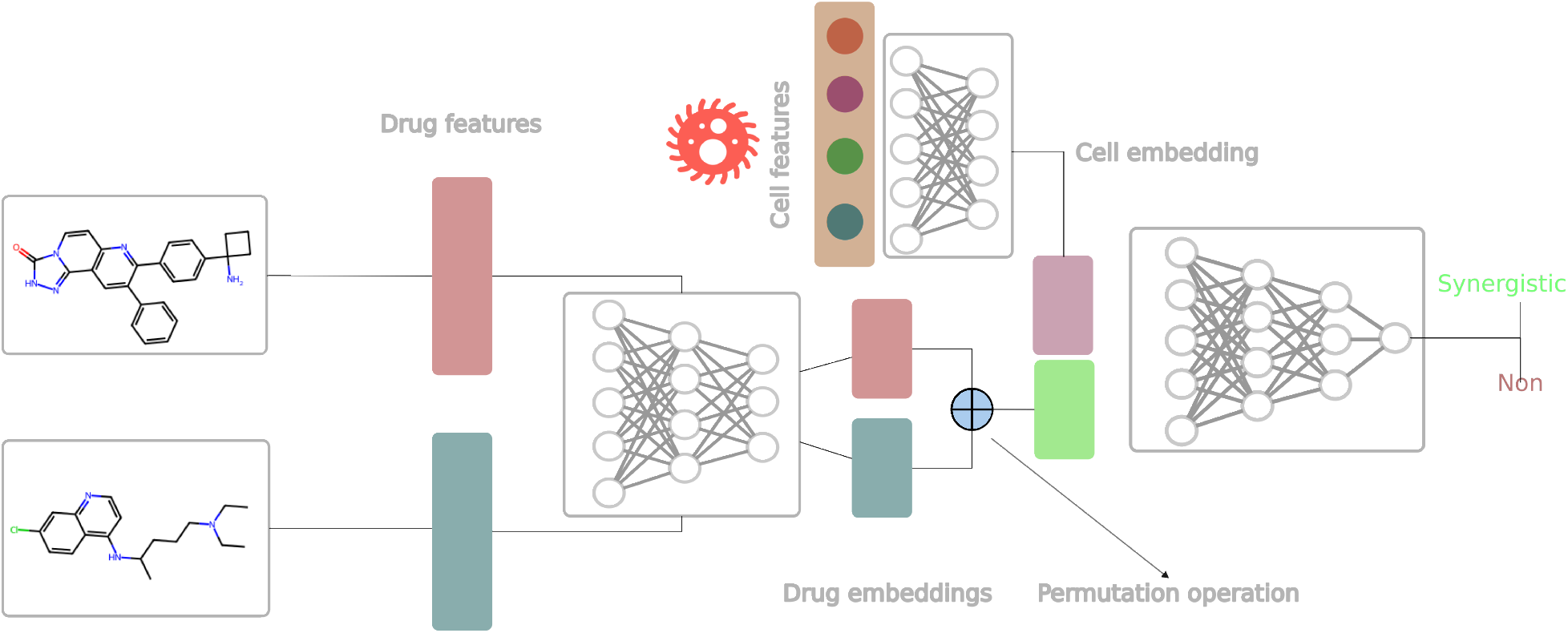
The MLP model for predicting synergy

**Figure 7.**
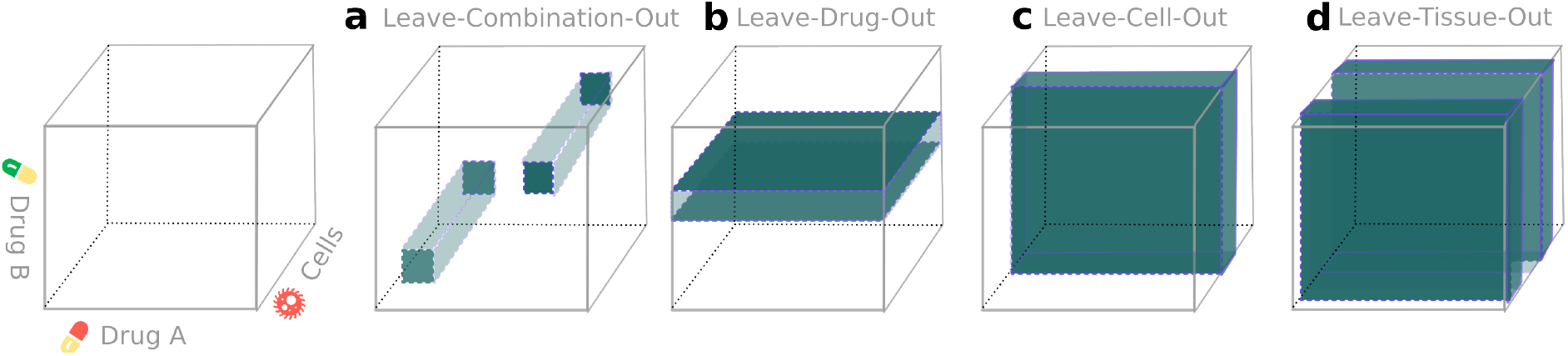
4 scenarios for splitting data to demonstrate the generalization ability of each algorithms. The overview of the (*DrugA, DrugB, cell*) combinatorial space is shown in a cube. (a) Leave-combination-out. (b) Leave-drug-out. (c) Leave-cell-out. (d) Leave-tissue-out.

Next, we investigated the batch size of the experimental platform, which greatly influenced the rate of synergy discovery (see Fig. 5b). 9 rounds were required for the manual platforms (*k* = 168) to identify 300 synergies, which correspond to only 1512 measurements. However, it requires 4476 measurements to discover the same amount of synergies when *k* = 896. Because our selection process enriches the batches with potential high synergies whereas increasing the batch size tends to introduce noise. As shown in Fig. SI5, when *k* = 168, among the top 168 predicted samples, the ratio of synergies is 0.36.

However, when *k* = 896, the ratio decreases to 0.19. More redundant samples are included with the increment of batch size. We observed the same behavior for the predictive qualities in PR-AUC as shown in Fig. SI6a. When the total number of training datasets reached 6127 samples, the PR-AUC can’t continue to improve even with more supplementary data. This tendency is even accentuated when we reduce *k* to 50 and 10 measurements, as shown in Fig. SI6b-c. We observed that using a smaller batch size led to the discovery of more synergistic samples. Additionally, based on PR-AUC, the exploration strategy outperformed the exploitation strategy during the initial rounds of sampling. When the training dataset size reached 300 data points, the exploitation strategy demonstrated a drastic improvement in predictions. Therefore, applying the hybrid strategy consistently identified more truly synergistic samples than the two other strategies (exploration and exploitation). This is because, in the early phase when very few batches were measured, identifying high-synergy drug combinations is difficult, confirmed by the poor performance of the exploitation criterion. In contrast, the exploration strategy is not affected by the lack of high-synergy samples in the training set. In our hybrid strategy, we switched from exploration to exploitation criterion as the increasing speed of PR-AUC slows down (see Materials and methods 6.7), which matches when the algorithm is capable of consistently predicting high synergies.

In conclusion, we confirm here that active learning has the potential to drastically improve the performances in synergy detection, reducing the experimental effort required for screening a combinatorial library of drugs. We showed that strategies are not equivalent: we observed that the best strategy is to start with the exploration criterion to quickly improve the predictive power of the MLP during the initial iterations and then switch to exploitation when the performance plateau is reached. Doing so, we evaluated that as few as 8 rounds of *k* = 168 is sufficient to extract 60% of the synergistic combination of *D* = 38 drugs across *C* = 29 cell types.

## 3 Discussion

While developing new drugs from scratch is very challenging and costly, drug repurposing with drug combinations offers an alternative for a potentially large array of diseases. However, identifying effective drug combinations is challenging due to the vast combinatorial space of drug pairs, from which only a small fraction are expected to be synergistic. Measuring combinations inhibition induced by drugs typically necessitates measurements across multiple concentrations of each drug. Although there have been vast drug combination screening campaigns that yielded the Almanac and Oneil datasets, these large-scale experimentations are typically not tractable. Therefore, AI has been introduced to use publicly available data to train synergy prediction algorithms that would recommend which combinations are likely to be synergistic. However, their performances are limited because the task is challenging—only a small fraction of the combinatorial space contains synergistic combinations. Recently, approaches based on active learning where the AI algorithm is refined iteratively by producing intermediate training data along the learning process have emerged. This approach has been shown to be very efficient [18, 25]. Here, we investigated the compositions of active learning procedures for predicting synergy in drug combinations by benchmarking various AI algorithms and active learning strategies. The goal of this work is to evaluate its components for one to build a specific active learning procedure for its own research.

In active learning, the AI algorithm for active learning is expected to operate at very low data regimes— being predictive with as few new measurements as possible. This depends on how the molecular information (molecular features) and cellular information (cellular features) are encoded to be fed to the algorithm. While the literature is rich in molecular features, all the ones we tested yielded similar results in prediction quality. In contrast, the cellular representation displayed a consistent gain. These results suggest that for synergy prediction, the importance of molecular properties (molecular topology for example) depends strongly on the cellular environment — the pathways influenced by one drug varies from one cell type to another. Therefore, improving the representation of the biological network at play in the cell is a promising avenue to improve synergy predictions.

For the AI algorithm itself, our results demonstrated that balancing the model complexity is crucial. Parameter-light algorithms such as logistic regression have a limited performance, whereas parameterheavy tend to have limited performance in low data regimes. We found that the MLP balanced the complexity of the model with the ease of tuning the hyper-parameters (depth and number of neurons). This suggests that even if larger NN with strong performance are largely available, they are difficult to adapt to new search spaces.

Indeed, these results are sensitive to the search space considered: in Oneil, we observed a saturation of the performances for all models whereas in Almanac, we did not. The latter suggest that the biological effects covered by the drugs in the combinatorial space of ALMANAC are vast. Therefore, the stage at which drugs are selected for the combinatorial search is critical.

Exploring a large combinatorial space involves making predictions across cell types or drugs for which no measurements or very few were made. In our benchmarks, when at least drugs and cells were measured in a few contexts is the most favorable case. In contrast, the prediction for unseen drugs is poor for all the tested algorithms (2 times lower than the combination). Indeed, the effect of a molecule can drastically vary depending on the pathways it affects, which may be triggered by very small molecular changes. For example, Lenalidomide is a derivative of Thalidomide with a minor structural change where an amino group replaces an oxygen atom in the phthalimide ring. But Thalidomide exerts its effect by inhibiting TNF-alpha, Lenalidomide influences the immunity cells more directly [32]. Across cell types, we showed that the algorithms generalize well; however, they did not across tissues. These results suggest that within the same tissue, drugs are affecting the same pathways, which are specific to the tissue. More generally, these results show the functional redundancy of cell viability as there are different pathways leading to cell inhibition.

Recent work has demonstrated the efficiency of active learning for drug pair synergy [18]; however, how to build the optimal framework remains challenging. We assembled several active learning strategies, which first reveal the importance of the batch size, where smaller and more frequent measurements drastically increase the synergy yield because smaller batches are richer in synergistic combinations—selecting of the best candidates only.

To select these candidates, the choices are based on one criteria that selects higher synergies (exploitation) or one that favors exploration in order to improve predictions quality (exploration). When the batch size is large, we did not observe striking differences between exploration-only or exploitation-only strategies. However, when the batch size decreases, the exploration yields a higher outcome than the exploitation in the first phase (*<* 300 measurements), the exploitation surpasses exploration. The mixed strategy did improve the detection power, where we essentially switched from exploration to exploitation when the AI algorithm converged in terms of prediction power. The rationale is that when the model has reached its maximum predictive capacity, no additional data will improve the detection rate. This observation generalizes to all AI algorithms beyond the MLP.

Finally, our recommendations are as follows:

- For the molecular features, molecular fingerprints offer sufficient information while being fast to compute and interpretable.
- For cellular features, gene expression profile, when available, carries critical information about the biological context.
- For the AI algorithm, MLP performs as well as other methods while being easy to adapt to specific applications and data sizes.
- For the selection strategy, the hybrid strategy is optimum by monitoring the performance of the algorithm along the cycles.

## 4 Availability of data and material

Code and data generated as well as fetch from other studies are available in the Github repository https://github.com/LBiophyEvo/DrugSynergy.git. The original data of drug combination and synergy scores can be downloaded from https://drugcomb.fimm.fi/jing/summary_v_1_5.csv. The cell genomic data can be downloaded from https://www.cancerrxgene.org/gdsc1000//GDSC1000_WebResources//Data/preprocessed/Cell_line_RMA_proc_basalExp.txt.zip.

## 5 Acknowledgements

We would like to thank Paul Dupuyds for useful discussion and code checking.

## 6 Materials and methods

### 6.1 Data collection and preprocessing

The data utilized in this study originates from O’Neil et al. [6]. This research investigated 39 cancer cell lines and 38 drugs, including 22 FDA-approved drugs. Comprehensive drug combinations involving FDA-approved drugs, as well as combinations of FDA-approved drugs with the remaining 16 non-approved drugs, were systematically tested, resulting in 583 drug pairs per cell line. We categorized the drug combination pairs using the Loewe synergy score, calculated via the DrugComb platform [5]. Following the recommendations from Abdullaev et al. [26], a threshold of 10 was selected for the classification task, as it is considered superior to a threshold of 5 for distinguishing between synergistic and non-synergistic drug pairs.

For independent validation, we utilized the NCI-ALMANAC database [7], which includes data on 60 cell lines and 5232 drug pairs tested across these lines, totaling 304,549 measurements. Detailed data statistics are presented in Table 2.

Our experimental setup required genomic features for each cell line; therefore, we retained only the data samples corresponding to cell lines included in the Genomics of Drug Sensitivity in Cancer (GDSC) dataset [29].

Cell line features were sourced from GDSC, which provides normalized gene expression profiles covering 17,737 genes across approximately 1,000 cancer cell lines. To reduce the dimensionality of the gene expression data, we selected 908 landmark genes as defined by the LINCS project [33], resulting in a 908-dimensional representation of each cell line.

Drug representations were derived from SMILES strings, and we evaluated five distinct molecular features: One-Hot-Encoding, Morgan fingerprints [19], MinHashed Atom-Pair fingerprints up to a diameter of four bonds (MAP4) [27], Molecular Access System (MACCS) [28], and representations pre-trained using ChemBERTa2 [12]. Morgan fingerprints with a radius of 2 and a length of 1024 bits and MACCS were generated using RDKit [34]. The MAP4 fingerprints, also with a radius of 2 and 1024 bits, were calculated using the MAP4 package [35].

### 6.2 Framework of MLP

MLP is a feed-forward neural network that outputs a single value: the probability of being synergistic or not. The architecture of MLP is depicted in Fig.6. The architecture consists of 4 modules:

1. Drug features dimension reduction layers: features for two drugs are fed into the same MLP to get their low-dimensional embedding representations (*h*(*D*_*i*_), *h*(*D*_*j*_)), respectively.

2. Cell features dimension reduction layers: these layers reduce the dimension of cell genomic features into a low-dimension representation, namely, cell embedding (*h*(*C*_*k*_)).

3. Permutation operation: this operation takes the embedding of drugs, transforming them into an intermediate embedding (*ĥ* (*D*)). This step is to ensure that the order of drugs won’t influence the output. Here we benchmarked three permutation invariant operations:

- Sum:

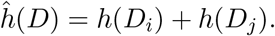
- Max:

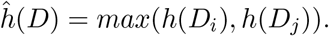
- Bilinear:

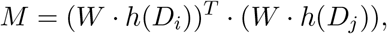

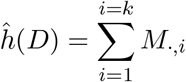

where *W* ∈ ℝ^*k×k×k*^ is a tensor, *k* is the dimension of the drug embedding *h*(*D*_*·*_). Each slice across the third dimension of *W* is a symmetric matrix.

4. Synergy prediction layers: this is an MLP whose input is the concatenation of cell embedding (*h*(*C*_*k*_)) and the intermediate embedding of drugs (*ĥ* (*D*)) after applying permutation operation. The output is the probability of being synergy.

The considered hyperparameters setting is summarized in Table.3. Cross Entropy was used as a loss function to be minimized.

### 6.3 State-of-art methods comparison

To evaluate data efficiency and accuracy, we employed the MLP described above along with five state-of-the-art methods. These methods include two simple machine learning models with light hyperparameter tuning, one medium-sized feed-forward neural network, and three advanced deep learning models that utilize graph neural networks or attention mechanisms with more complex hyperparameters.

- Logistic Regression (LR): LR assumes that the input variables are linearly related to the output. In classification, the output is finalized by using a logistic function. LR was used to compare the other models with a linear method.
- XGBoost [36]: An ensembling learning method based on the decision tree and gradient descent. Compared with the traditional gradient descent method, XGBoost is less prone to overfitting. Compared with the representative decision tree model such as random forest, XGBoost is more accurate and more flexible.
- DeepDDS GCN and DeepDDS GAT [13]: The features of drugs are obtained by using a Graph Attention Network (GAT) or Graph Convolutional Network (GCN). In the graph, each node represents each atom of the drug molecule, it is initialized by a set of atomic attributes. The edge of the graph shows the chemical bond between atoms. The cell line features are obtained by using 954 landmark genes (based on LINCS project [37]) from Cancer Cell Line Encyclopedia (CCLE) [38]. Firstly, an MLP is used to extract the cell line feature. Then it is concatenated to the drug features obtained by GCN or GAT as an input for the MLP-based classifier.
- DTSyn: Similarly to DeepDDS GCN, GCN blocks concatenated with gene embeddings are put through a fine-granularity transformer encoder block in order to extract chemical substructures–gene interactions. Meanwhile, gene embedding and GCN-obtained drugs’ features are fed into a coarse granularity transformer encoder block. The high-level features obtained from these two transformers are used to predict synergy labels by an MLP.

The comparison of the number of parameters for NN based model is in Table.4:

### 6.4 Metrics

We employed Area Under the Precision-Recall Curve (PR-AUC). This metric can display how well the model can identify the positive samples. It is more suitable for the unbalanced dataset. Since our model is highly unbalanced, and the objective is to explore more positive samples, PR-AUC will be a much more important indicator. For a random classifier, the PR-AUC will be the ratio of positive samples in the dataset.

In Section 2.2, we also utilized recall, defined as the proportion of true positive samples among all positive samples. Recall measures the fraction of relevant targets that the algorithm successfully retrieves, indicating its capability to identify new candidates.

### 6.5 Data split scenarios

In order to check if the model can successfully predict the new drug combinations, new cell lines, and new drugs. The following scenarios are designed as shown in Fig.7. The drug combinations are actually triple combinations of (*DrugA, DrugB, cell*). The 4 scenarios are illustrated in the following:

- Leave-combination-out scenario: this scenario is used to mimic the situation where the biologist needs to test new drug combinations based on the drugs and cells that have already been tested in other contexts. In the experiments, 50-cross-validation is used to assess the model performance.
- Leave-drug-out scenario: this aims to simulate the case where some new anti-tumor agents are expected to be tested. Each distinct drug within the dataset underwent partitioning into *Train drug* and *Test drug* sets through 20-fold cross-validation. Afterward, all samples with either DrugA or DrugB in the *Test drug* set will be put aside as the test dataset.
- Leave-cell-out scenario: this procedure is implemented to assess the predictive capacity of the model for novel cell lines. Within each experimental iteration, a single cell is designated as *Test cell*, following which all drug combinations tested on this specific *Test cell* are allocated to form the test dataset. Thus, the data partitioning strategy in this context adheres to a *N*_*cell*_ cross-validation, where *N*_*cell*_ represents the total number of cells encompassed within the dataset.
- Leave-tissue-out scenario: this experimental scenario seeks to evaluate the model’s ability to predict synergy across novel cell lines originating from distinct tissue types. This is a notably more challenging task than the previous assessment. In each iteration, a specific tissue is designated as *Test tissue*, with all measurements pertaining to this tissue being reserved for the test dataset. A *N*_*tissue*_ cross-validation approach was adopted, where *N*_*tissue*_ corresponds to the number of tissue types present in the dataset. The Oneil dataset comprises cells sourced from six distinct tissues, the Almanac dataset includes cells originating from nine different tissues, thus expanding the diversity of cell line origins (See Table.2).

### 6.6 Number of samples per round in active learning

To align the active learning process more closely with practical scenarios, we opted for a sample size in each iteration to approximate that of experimental conditions.

As for Oneil dataset [6], they used a robotic platform for conducting the experiments. Cells are plated in 1536-well plates. Besides, cells are treated with a 4 by 4 matrix of drug concentration and there are 4 replicates for each drug combination concentration. The incubation time is 4 days. As for Dream dataset [9], they used 384 well plates, and for every plate, and tested 4 combinations per plate in a 6 × 6 format [39]. Each drug combination is in 5 replicates. The incubation time is 5 days. Since the drug screening data of Almanac was collected from three different institutions, the protocol changes a little bit from each other, so we mainly consider the experimental scenarios in Oneil and Dream, which represent an automatic platform and a manual platform respectively.

In a conventional incubator, it has the capacity to incubate either 56 units of 1536-well plates or 42 units of 96 or 384-well plates. In laboratories lacking automated equipment, the typical choice is the 384-well plates. Assuming four-drug combinations are to be executed on a plate similar to the Dream dataset, and with the availability of one incubator, a parallel execution of 168 drug combinations becomes feasible. However, employing a robotic system enables biologists to utilize 1536-well plates, resulting in a drug screening capacity four times greater than that of the 384-well plates, accommodating 16 drug combinations per plate. With one incubator at disposal, parallel execution facilitates a throughput of 896 drug combinations. Therefore, in our experiments, we adopted 168 and 896 drug combinations per round to emulate the two different experimental setups depending on the laboratory facilities.

Furthermore, the typical throughput was five cell lines per day by automatic platform [6]. So We restricted that no more than 5 cell lines will be added in each round.

### 6.7 Candidate selection strategies in active learning

The idea is to systematically choose the cell lines and corresponding drug combinations that are expected to provide the most reliable information, thus improving the overall confidence in experimental outcomes. Here, each round involves a total of *k* experimental slots, dividing them evenly among the top 5 cell lines ensures a balanced and targeted approach.

By using an ensemble approach [31], each AI algorithm are trained *M* times simultaneously. Therefore for each inquired sample, AI algorithm will give a *M* instances of prediction: *S*_1_, *S*_2_, *…, S*_*M*_. This ensemble method provides us with the mean and uncertainty (*S, σ*(*S*) of prediction for each candidate triplet.

Three selection criteria are employed to optimize the synergy discovery based on (*S, σ*(*S*).

- Exploration: This strategy prioritizes candidates that are expected to enhance the confidence of the predictions by selecting samples with high *f* (*S*) = *σ*(*S*), as these are likely to reduce uncertainty in the results.
- Exploitation: This approach focuses on selecting samples that are likely to maximize the objective, which in this context is identifying more synergies. Therefore, samples with high *f* (*S*) = *S* values are preferred, as they are expected to exhibit stronger synergistic effects.
- Hybrid: This strategy aims to balance exploration and exploitation throughout the learning process. In the early phases (within the first 3 rounds), high-synergy data often have high uncertainty. As the active learning progresses, the uncertainty associated with high-synergy samples decreases, as illustrated in Fig. SI5. Relying on a single strategy, either exploration or exploitation, can lead to sub-optimal outcomes. To effectively switch between exploration and exploitation, we employ an acquisition function defined as *f* (*S*) = *α* ·*S* + (1 −*α*) · *σ*(*S*). Here, *α* controls the balance between focusing on high-synergy samples *S* and samples with high uncertainty *σ*(*S*). Here we used a :

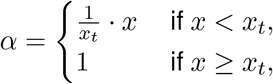

where *x* represents the derivative of PR-AUC (Precision-Recall Area Under the Curve), and *x*_*t*_ denotes the transition point where the rate of increase in PR-AUC begins to slow down. When PR-AUC slows down, it indicates that adding more data yields diminishing returns in improving the algorithm’s performance, suggesting that the model is already relatively accurate. At this point, it is advantageous to shift towards exploitation by increasing the weight *α* on *S* in the acquisition function, thus prioritizing high-synergy samples over those with high uncertainty.

Besides, since the traditional throughput of cells is 5, we constraint that, in each round, no more than 5 cell lines will be selected. Here is the breakdown of the selection process:

1. Selection of Cell Lines:
  - In order to identify the top 5 cell lines that are likely to improve the confidence of predictions, we compute the average value of *f* (*S*) for each cell line.
  - Rank the cell lines based on their averaged *f* (*S*) values and choose the top 5 cell lines with the highest averages.
2. Ranking and Selection of Drug Combinations:
  - For each selected cell line, rank the available drug combinations by their *f* (*S*) value.
  - From these rankings, select the top *k/*5 drug combinations for each cell line. This means that for a total of *k* experimental candidates, each of the 5 selected cell lines will contribute *k/*5 drug combinations evenly.

### Contributions

AA, PN, and VO conceptualized the project; SW and VO performed the research and wrote the original draft; and all authors reviewed the manuscript.

