## Supplementary material for "A Guide for Active Learning in Synergistic Drug Discovery": drug_synergy_prediction_si.pdf

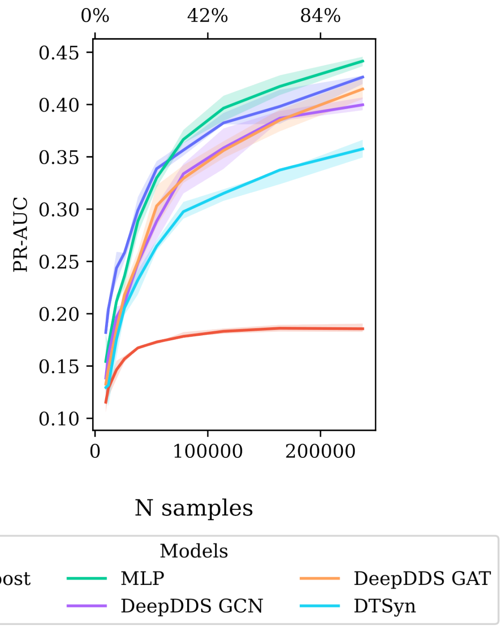


Fig. SI1: Performance of algorithms with varying proportions of training data on Almanac dataset. The X-axis shows the number of training data points, and the Y-axis displays the PR-AUC on the validation dataset. Solid lines represent average performance over 10 random cross-validations, with shaded areas indicating the 0.9 quantile range.

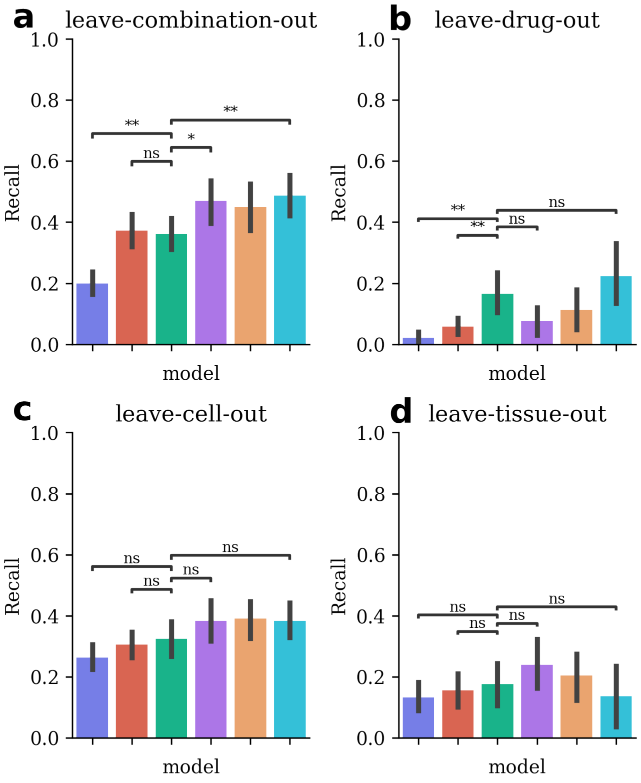


Fig. SI2: The Recall of distinct models on each scenario demonstrates the generalization ability of algorithms. Each scenario corresponds to a different way of splitting the data, displaying different generalization perspectives of each algorithm. (a) Leave combination out is random split of data. (b) leave drug out. 5% of drugs are excluded. (c) Leave cell out. Every time, all drug combinations on one specific cell are excluded as validation. (d) Leave tissue out. All cells from one tissue are excluded as validation. X-axis shows all models, the Y-axis is the Recall value. ns: $0.05 < p \leq 1$, $* : 0.01 < p \leq 0.05$, $** : p <= 0.01$. Algorithms are evaluated on each task by using cross-validation.

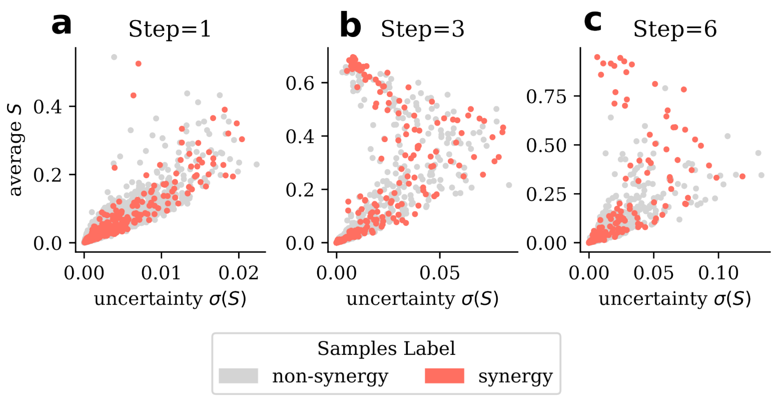


Fig. SI3: Mean versus standard deviation of predicted probabilities from $M$ ensembled MLPs in the 1st, 3rd, and 6th rounds of active learning with batch size $k = 168$. Pink dots represent non-synergistic samples, and gray dots represent synergistic samples. (a) 1st Iteration: Standard deviation and mean of each sample are highly correlated. (b) 3rd Iteration: High-synergy samples begin to shift towards the low-uncertainty regime. (c) 6th Iteration: Most high-synergy samples cluster around the low-uncertainty regime. Initially, the standard deviation and mean of predictions are strongly correlated, showing little difference between exploration and exploitation. However, after several iterations, the model becomes more accurate in predicting high synergy, with high-synergy samples increasingly appearing in the low standard deviation regime.

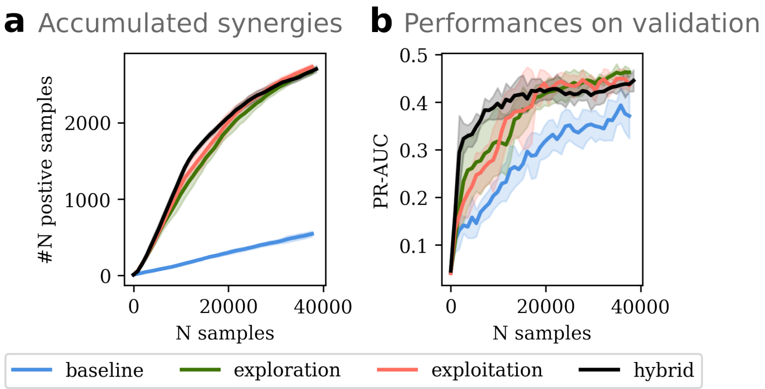


Fig. SI4: Application of active learning on the Almanac dataset, comparing exploration (green), exploitation (pink), and hybrid (black) strategies with batch size $k = 896$. (a) Accumulated number of synergistic samples over successive experimental rounds. (b) Performance of the NN algorithm as more data is sequentially incorporated. This comparison highlights how different strategies influence synergy discovery and algorithm performance throughout the active learning process. The active learning strategy also works on big datasets.

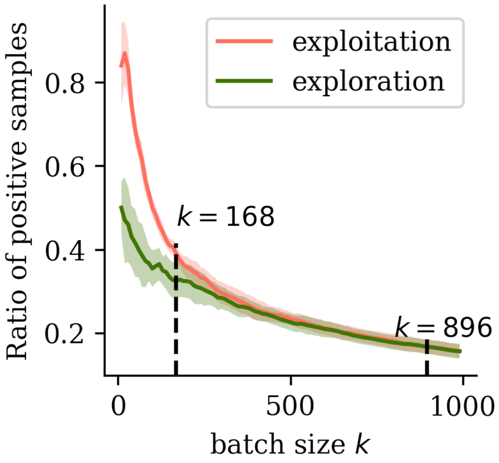


Fig. SI5: Ratio of synergies among the top $k$ predicted samples, comparing exploration and exploitation strategies. As $k$ increases, the ratio of identified synergistic targets decreases, highlighting the diminishing return on synergy discovery with larger batch sizes.

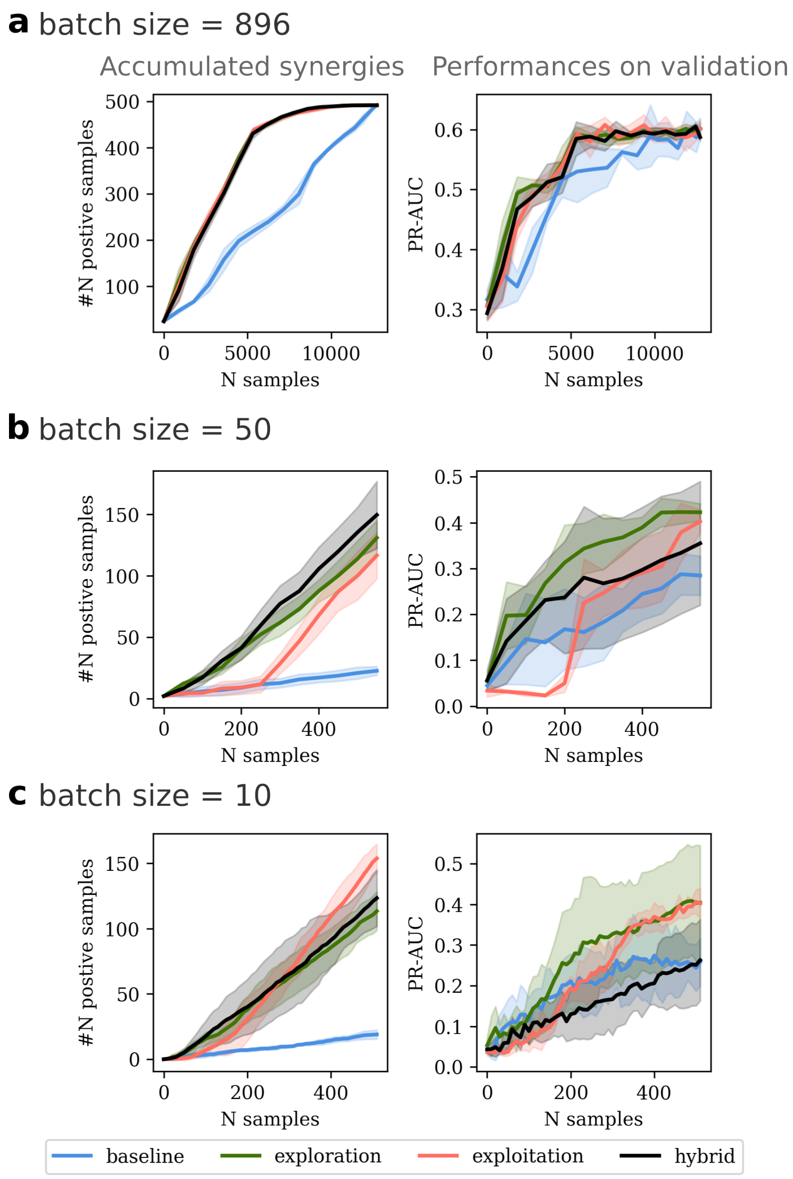


Fig. SI6: Comparison of synergy yield and algorithm performance across varying batch sizes and selection strategies, including exploration, exploitation, and hybrid approaches. Left Panels: Accumulated number of synergies as more data is sequentially added. Right Panels: PR-AUC performance with iterative data addition. (a) Batch size $k = 896$. (b) Batch size $k = 50$. (c) Batch size $k = 10$.
